# Translation enhancement by a *Dictyostelium* gene sequence in *Escherichia coli*

**DOI:** 10.1101/564815

**Authors:** Tomo Kondo, Shigehiko Yumura

## Abstract

Methods for heterologous protein production in *Escherichia coli* have revolutionized biotechnology and the bioindustry. It is ultimately important to increase the amount of protein product from bacteria. To this end, a variety of tools, such as effective promoters, have been developed. Here, we present a versatile molecular tool based on a phenomenon termed “translation enhancement by a *Dictyostelium* gene sequence” (“TED”) in *E. coli*. We found that protein expression was increased when a gene sequence of *Dictyostelium discoideum* was placed upstream of the Shine-Dalgarno sequence located between the promoter and the initiation codon of a target gene. The most effective sequence among the genes examined was *mlcR*, which encodes the myosin regulatory light chain, a subunit of myosin II. Serial deletion analysis revealed that at least 10 bases of the 3’ end of the *mlcR* gene enhanced the production of green fluorescent protein in cells. We applied this tool to a T7 expression system and found that the expression level of the proteins tested was increased when compared with the conventional method. Thus, current protein production systems can be improved by combination with TED.

## Introduction

*Escherichia coli* is one of the most powerful workhorses in biological science, including biotechnology, because of its low cost, tractability, and the availability of conventional methods (Rosenberg et al. 1987; Qing et al. 2004; Structural Genomics Consortium et al. 2008). Over the last four decades, a great variety of proteins, including somatostatin (Itakura et al. 1977), insulin (Goeddel et al. 1979), interferon (Taniguchi et al. 1980; Derynck et al. 1980), interleukin (Devos et al. 1983), and Cas9 (Jinek et al. 2012; Gasiunas et al. 2012), have been bacterially synthesized in the laboratory and at the industrial scale. More recently, bacterial synthesis of antibody fragments (e.g., Fab and nanobody) has been reported to minimize the cost of and the sacrifice of animals for production (Spadiut et al. 2014; Buser et al. 2018). Thus, protein production using genetically engineered *E. coli* benefits the understanding of fundamental cellular processes as well as the development of biopharmaceuticals.

To increase protein yield, it is effective to increase the absolute amount of protein production in the cell. The first approach is to increase the transcription of the target gene by using strong promoters, such as *lac* and its derivatives (Dickson et al. 1975; de Boer et al. 1983). In addition, expression levels can be regulated by three factors of the sequence upstream of the initiation codon of the target gene: (i) the strength of the Shine-Dalgarno (SD) sequence (Shine and Dalgarno 1974), (ii) the distance between the SD sequence and the initiation codon (Singer et al. 1981; Shepard et al. 1982; Ringquist et al. 1992; Chen et al. 1994a), and (iii) the sequence upstream of the SD sequence (Roberts et al. 1979; Stanssens et al. 1985; Coleman et al. 1985; McCarthy et al. 1985; Olins et al. 1988). In factors (i) and (ii), optimum patterns can be determined by a complementary sequence of the 3’ end region of 16S rRNA and the relative position of the 30S ribosome, respectively. The third factor is known to affect mRNA stability and the preferred interaction of mRNA with the ribosomal S1 protein (Boni et al. 1991; Komarova et al. 2002; Komarova et al. 2005; Takahashi et al. 2013). According to these studies, an A/U-rich sequence, with typically more than 70% AT content at the 5’-untranslated region (UTR), is empirically effective, but which sequences can efficiently induce protein expression remains elusive.

Here, we report that gene sequences from the social amoeba *Dictyostelium discoideum*, which has an A/T-rich genome (Eichinger et al. 2005), enhance expression in *E. coli*. We tested genes having 60–73% AT content and found that even a sequence an AT content of less than 70% has the ability to enhance expression. The most effective sequence was the *mlcR* gene, which encodes the myosin regulatory light chain, an evolutionarily conserved subunit protein that regulates the function of myosin II (Uyeda and Spudich 1993; Chen et al. 1994b; Liu et al. 1998; Kondo et al. 2011, 2012, 2015). As proof-of-principle, we demonstrate the usefulness of this sequence for expression enhancement by assessing the expression of proteins, including green fluorescent protein (GFP), in *E. coli*.

## Materials and methods

### E. coli culture

*E. coli* strains HST08 Premium (Takara Bio, Shiga, Japan), DH5α (Cosmo Bio, Tokyo, Japan), and BL21(DE3) (Cosmo Bio) were used. The cells were cultured at 37°C in LB medium (Nippon Genetics, Tokyo, Japan) supplemented with 100 µg/ml ampicillin (Wako Pure Chemical Industries, Osaka, Japan) or 20 µg/ml kanamycin (Wako Pure Chemical Industries). Luria-Bertani (LB) agar was prepared by solidifying LB medium with 1.5% agar (Wako Pure Chemical Industries). For expression induction by isopropyl β-D-thiogalactopyranoside (IPTG) (Nacalai Tesque, Kyoto, Japan), 1/100 volume of precultured cells were inoculated in fresh LB medium containing appropriate antibiotics and were cultured at 37°C until an optical density at 600 nm of 0.5–0.6 was reached. Then, IPTG was added at the final concentration of 0.5 mM, followed by culture at 22°C for 16 h or at 37°C for 3 h.

### Plasmids and cDNA cloning

Isolation of plasmid DNA, preparation of DNA fragments, ligation, and transformation of *E. coli* cells were carried out using standard techniques. The genes of *D. discoideum* were amplified from cDNA (Robinson and Spudich 2000) or genomic DNA of the Ax2 strain using PrimeStar Max DNA polymerase (Takara Bio) and the designed primers, and were cloned into pUC19 (Yanisch-Perron et al. 1985) (Takara Bio). The cDNA was kindly provided by Dr. Douglas Robinson (Johns Hopkins University School of Medicine, Baltimore, MD, USA). Genomic DNA was purified from Ax2 cells with the NucleoSpin Tissue kit (Takara Bio). The following plasmids were gifted by Dr. Scott Gradia: pET His6 GST TEV LIC cloning vector (1G) (Addgene plasmid # 29655), pET His6 MBP TEV LIC cloning vector (2M-T) (Addgene plasmid # 29708), pET His6 Sumo TEV LIC cloning vector (2S-T) (Addgene plasmid # 29711), and pET GFP LIC cloning vector (2GFP-T) (Addgene plasmid # 29716). The reverse complementary sequence of *mlcR* was synthesized by Integrated DNA Technologies, Inc. (Coralville, IA, USA).

### Imaging and quantification

For excitation of GFP in bacteria grown on an agar plate, a custom-made blue light-emitting diode (LED) illuminator was used. For imaging of *E. coli*, culture was dropped on a coverslip (24 × 60 mm; Matsunami Glass, Osaka, Japan) and overlaid with a block of 1.5% agar (8 × 8 × 1 mm; Dojindo Laboratories, Kumamoto, Japan) in distilled water. The specimens were observed with an inverted microscope (DIAPHOT300; Nikon, Tokyo, Japan) equipped with a camera (Orca-ER C4742-80-12AG; Hamamatsu Photonics, Hamamatsu, Japan) and a mercury lamp (Nikon). Images were acquired five times using a 100× objective lens (Nikon), and maximum intensity projection was performed using Fiji (Schindelin et al. 2012). To obtain a Z score, the cytoplasmic fluorescence intensity was normalized by subtracting the mean intensity from 10 background regions, and the value was divided by the standard deviation of the intensity of the background regions. The statistical significance of differences between groups for a dataset was tested by the Wilcoxon rank sum test (“ranksum” function in MATLAB; MathWorks, Inc., Natick, MA, USA).

### Prediction of RNA secondary structure

The local secondary structure of RNA was predicted using RNAfold (Gruber et al. 2008) with default settings. Centroid structures, which are the secondary structures with minimal base pair distance to all other secondary structures in the Boltzmann ensemble, are shown in relevant figures.

## Results

### Protein expression in *E. coli* is enhanced by *Dictyostelium* gene sequences

We cloned the *mlcR* gene from a cDNA library of *D. discoideum* Ax2 and linked it with *gfp* in the pUC19 vector (Fig. 1a). These two genes were connected with nucleotides composed of an SD sequence-spacer (Fig. S1). After transformation of the plasmid into *E. coli* (strain HST08) and overnight culture on LB agar in the dark, we found that the colonies showed fluorescence upon excitation with blue LED light, even in the absence of the expression inducer IPTG (Fig. 1b, bottom-right). In contrast, no visible fluorescence appeared in cells harboring the plasmid containing only the SD sequence-spacer and *gfp* (Fig. 1b, top-left). GFP expression did not adversely affect the growth rate (Fig. S2). A similar level of fluorescence was detected in another strain, DH5α (Fig. S3). Moreover, fluorescence could be observed in cells expressing not only GFP, but also the red fluorescence protein mRuby3 (Bajar et al. 2016) (Fig. 1c). Microscopic observation revealed that fluorescence was observed in the cytoplasm of individual cells (Figs. 1d and e). Thus, the *mlcR* sequence enhances the expression of the downstream gene.

**Fig. 1.**
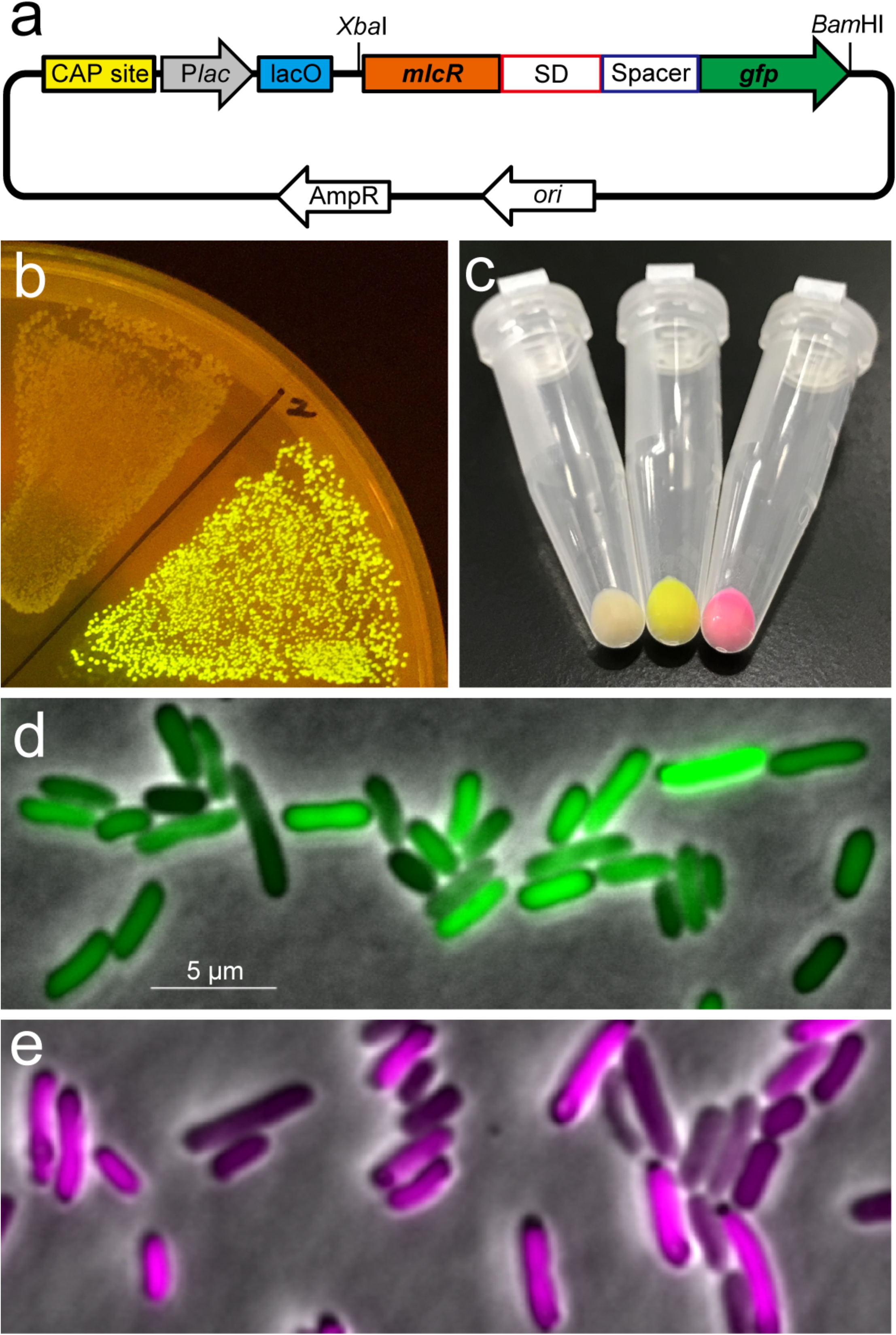
GFP expression in *E. coli* harboring the plasmid containing an *mlcR* gene sequence upstream of the SD sequence, a spacer, and *gfp*. (a) *mlcR*–*gfp* was cloned between the *Xba*I and *Bam*HI sites in the pUC19 vector. The nucleotide sequence of the insert is shown in Fig. S1. The SD sequence is designated as “SD.” (b) GFP fluorescence of *E. coli* (strain HST08) on an LB-agar plate not containing IPTG. The bacteria harboring the plasmid described in (a) in the bottom right of the figure exhibit GFP fluorescence. The upper left bacteria without fluorescence contain a similar plasmid lacking the *mlcR* gene. The plate was photographed over a UV shield during irradiation with a hand-held blue-light LED illuminator. (c) Bacterial pellets with or without fluorescence. Left; no fluorescence, center; GFP fluorescence, right; mRuby3 fluorescence. (d and e) Merged fluorescence and phase-contrast microscopic images of cells expressing GFP (green) or mRuby3 (magenta).

Next, we investigated whether gene sequences other than *mlcR* (68% AT content) also induce expression enhancement. Previous studies have reported translation enhancement, which is induced by A/U-rich sequences upstream of the SD sequence (McCarthy et al. 1985; Komarova et al. 2002). Thus, we additionally tested arbitrarily selected 8 genes, including *H1, eb1, tubC, tom7, tom20*, DDB_G0288629, *tom40*, and *tom70*, having AT contents of 60–73% (Fig. 2a, Table S1). All of the genes tested showed expression enhancement activity of various strengths (Fig. 2b). Hereafter, we refer to this phenomenon as “TED” (“translation <u>e</u>nhancement by a *<u>D</u>ictyostelium* gene sequence”).

**Fig. 2.**
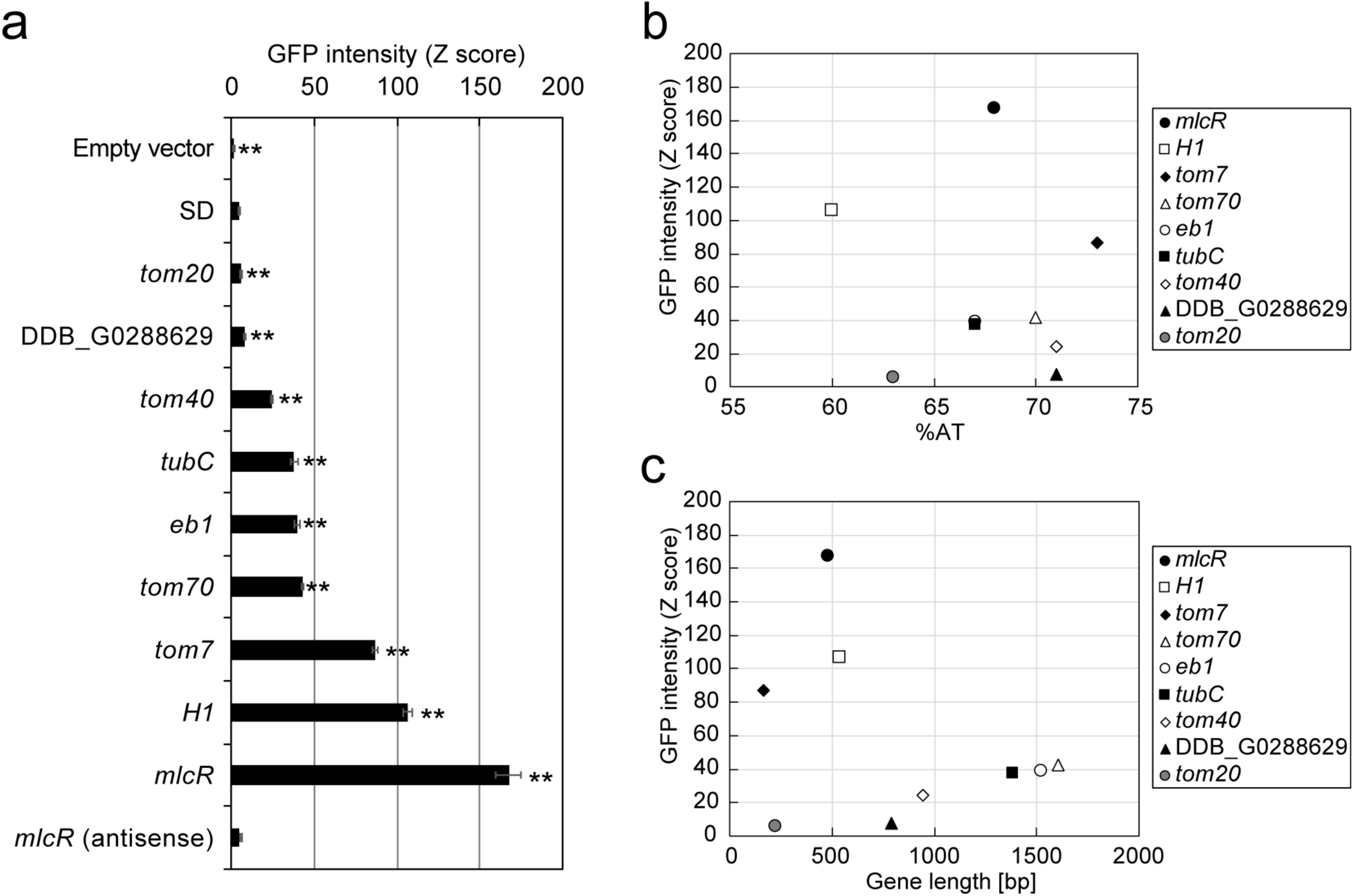
Enhancement of GFP expression in *E. coli* harboring plasmid containing the full length of the indicated gene sequence. (a) The strength of induction of GFP expression differed depending on the gene. All genes were inserted into the pUC19 vector, as shown in Fig. 1A. As a negative control, cells harboring empty pUC19 vector and only SD sequence, designated as “SD,” were tested. GFP expression can be visually confirmed with a Z score of approximately 80. Error bars represent the standard errors of the means. All statistical tests are the result of comparison with SD. n ≥ 239. \*\**p* < 0.01 (Wilcoxon rank sum test). (b) Scatter plot of GFP intensity versus AT content of the indicated gene. (c) Scatter plot of GFP intensity versus nucleotide length of the indicated gene.

The *mlcR* gene (68% AT content, 483 bp) had the highest TED activity among the genes examined. TED activity was measured as the ratio increase in GFP fluorescence. As shown in Figs. 2b and c, there was no obvious association between the strength of TED ability and the AT content or the length of the gene. The *H1* gene (60% AT content, 540 bp) and the *tom7* gene (73% AT content, 165 bp) exhibited relatively high TED activity. In contrast, some of the genes that had an AT content of more than 70% (i.e., DDB_G0288629 [71%, 789 bp], *tom40* [71%, 942 bp], and *tom70* [70%, 1605 bp]) or were similar to *mlcR* in AT content (i.e., *eb1* [67%, 1518 bp] and *tubC* [67%, 1386 bp]) showed relatively weak activity. These data suggest that TED activity cannot be simply explained by the simple parameters AT content and gene length. Consistent herewith, the complementary sequence of *mlcR* no longer had a translation enhancement effect (Fig. 2a). Thus, the simple concept of using an AU-rich sequence for enhancing expression was proven insufficient.

### TED activity can be achieved with a 3’ end of at least 10 bp of the *mlcR* sequence

Hereafter, we focus on *mlcR*, which had the strongest TED activity. To analyze the mechanism of expression enhancement, we identified the sequence that is indispensable for enhanced protein expression by *mlcR* deletion analysis. As the inducer agent (i.e., IPTG) was not added to the medium, we hypothesized that the basal activity of the *lac* promoter with the catabolite activator protein (CAP) site contributes to the expression of *gfp*, as previously described (Kennell and Riezman 1977; Yu and Reznikoff 1984; Bellis and Schwartz 1990; Wilson et al. 2007; Gatti-Lafranconi et al. 2013). As expected, GFP fluorescence disappeared for all examined patterns that lacked both the *lac* promoter (P*lac*) and the *lac* operator (lacO), only P*lac*, or the CAP site (Fig. 3a), suggesting that P*lac*-dependent transcription, not a potential cryptic promoter in the *mlcR* sequence, is responsible for *gfp* expression.

**Fig. 3.**
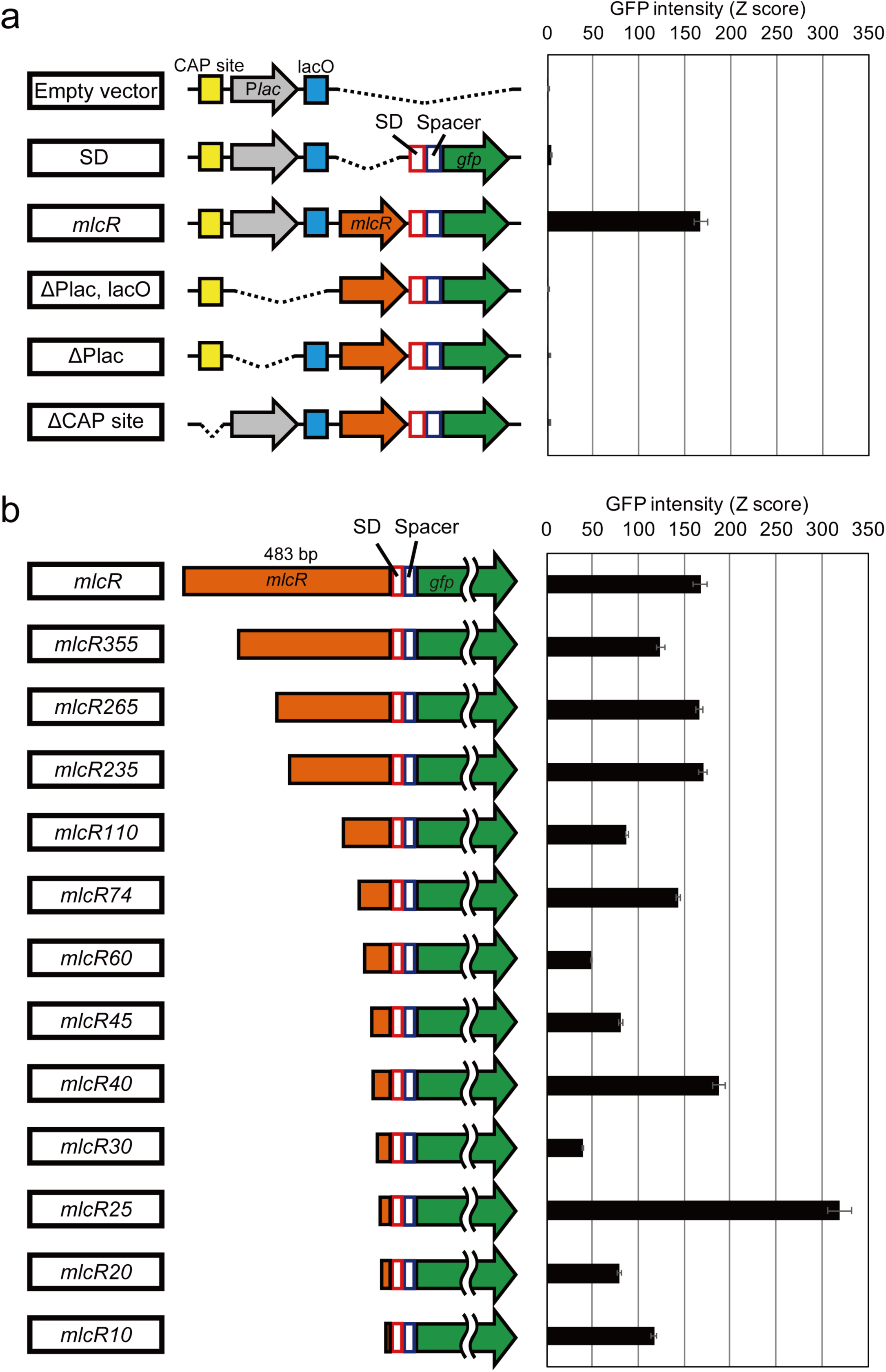
Analysis of the region required for GFP expression in the plasmid containing *mlcR*–*gfp*. (a) Deletion of the *lac* promoter (P*lac*), the lac operator (*lacO*), and/or the CAP site reduced GFP expression. Error bars represent the standard errors of the means. n ≥ 239. (b) Serial deletion of the *mlcR* sequence in the pUC19 vector. The number that follows “*mlcR*” is the number of bases from the 3’ side that does not contain the stop codon of the *mlcR*. The value of *mlcR* is the same as that shown in (a). Error bars represent the standard errors of the means. n ≥ 344.

Next, we investigated which region of the *mlcR* sequence is required for GFP expression. The sequence was deleted serially from the 5’ end of full-length *mlcR* cDNA (483 bp) (Figs. 3b and S1). In the *mlcR* sequence, there are eight potential initiation codons (ATG) in the frame that can express *gfp*. However, there are no sites with an SD sequence (e.g., AGGA, GAGG, or GGAG) upstream (at least 15 bp) of the initiation codon. Thus, the SD sequence downstream of the *mlcR* sequence is the only sequence to express *gfp*. We found that even only 10 bp of the 3’ end of *mlcR* (*mlcR10*) had TED ability. The 25-bp fragment of the 3’ end of *mlcR* (*mlcR25*) showed the highest activity. Interestingly, the 5-bp longer construct (*mlcR30*) showed the lowest activity. These data suggest that TED activity can be controlled by using at least 10 bp or longer of the *mlcR* sequence.

### Predicted local secondary structure around the ribosome-binding site

The secondary structure of RNA affects the translation rate (Iserentant and Fiers 1980; Gold et al. 1981; de Smit and Duin 1990; de Smit and van Duin 1990; Osterman et al. 2013). To gain insight into the variation in strength of TED activity using parts of the *mlcR* sequence of various lengths, we examined the predicted local structure of RNA. To focus on the property of the inserted sequence, the inserted *mlcR* sequence and the initiation codon were used for calculation. The sequences with relatively high expression upregulation (*mlcR10* [Fig. 4a], *mlcR20* [Fig. 4b], *mlcR25* [Fig. 4c], *mlcR40* [Fig. 4e], *mlcR45* [Fig. 4f], and *mlcR74* [Fig. 4h]) were expected to have a stem-loop structure near the initiation codon, whereas such structure was lost in *mlcR30* (Fig. 4d) and *mlcR60* (Fig. 4g), which had relatively low TED activity. These correlations suggest that the property of forming a stem loop consisting of an SD sequence and a proper length of the *mlcR* sequence is suitable for effective translation.

**Fig. 4.**
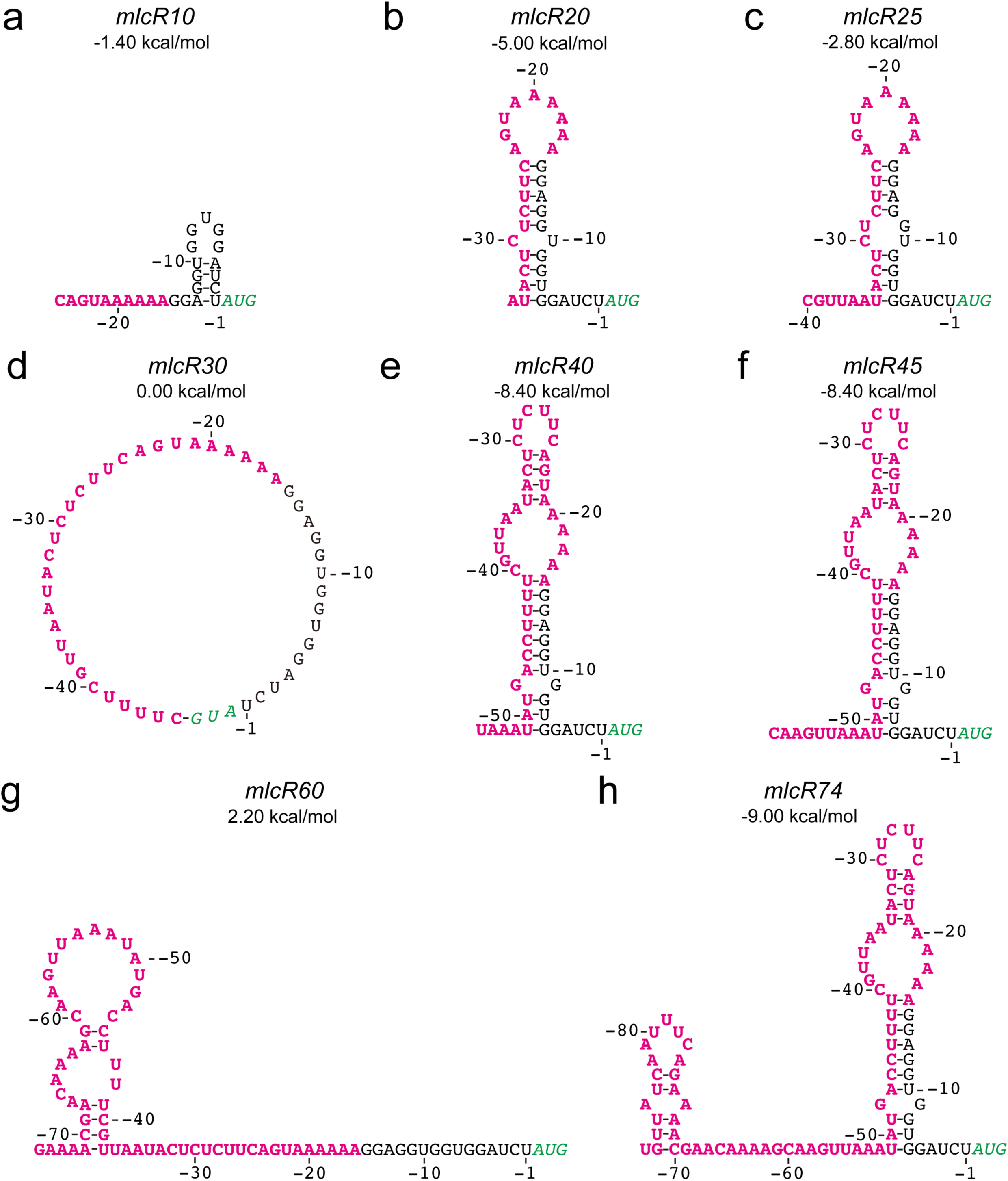
mlcR sequences with relatively high TED activity form a stem loop with the SD sequence. Predicted secondary RNA structures of *mlcR*10 (a), *mlcR*20 (b), *mlcR*25 (c), *mlcR*30 (d), *mlcR*40 (e), *mlcR*45 (f), *mlcR*60 (g), and *mlcR*74 (h). The mlcR sequence and initiation codon are shown in magenta bold and green italics, respectively. The calculated minimum free energies at 37°C are indicated.

### *mlcR25* TED improves the expression level of the classical pET expression system

We combined TED with the pET expression system (Studier et al. 1990). The pET vectors contain the T7 phi10 promoter and related sequences upstream of the start codon of a target protein. To demonstrate the effect of TED, we replaced 22 bp of the sequence between the SD sequence and an *Xba*I site with *mlcR25* (Fig. 5a). The similarity of these sequences was 41% (9/22 nucleotides) at most. According to the prediction of the local RNA structure, both the T7 phi10 sequence (pET-native) and *mlcR25* formed a similar stem-loop structure (Fig. 5b and c). Using these vectors, we expressed several types of proteins in *E. coli* strain BL21(DE3), which is frequently used for high-yield production. The formation of protein aggregates (i.e., inclusion bodies) is generally prevented at low temperature (Schein and Noteborn 1988); therefore, low-temperature culture is usually used to increase yield (Structural Genomics Consortium et al. 2008). Thus, we tested two culture temperatures, 22°C and 37°C. After IPTG induction, GFP (Fig. 5d), GFP-tagged maltose binding protein (MBP) (Fig. 5e), small ubiquitin-like modifier (SUMO) (Fig. 5f), and glutathione-S-transferase (GST) (Fig. 5g) were successfully expressed in cells harboring *mlcR25*-inserted plasmid, and the fluorescence level was increased by approximately 10% compared to that in cells harboring the original T7 phi10-inserted plasmid, at 22°C. A similar effect was observed in cells cultured at 37°C (Fig. 5h-k). These data suggest that the classical pET expression system can be improved by using *mlcR25* TED.

**Fig. 5.**
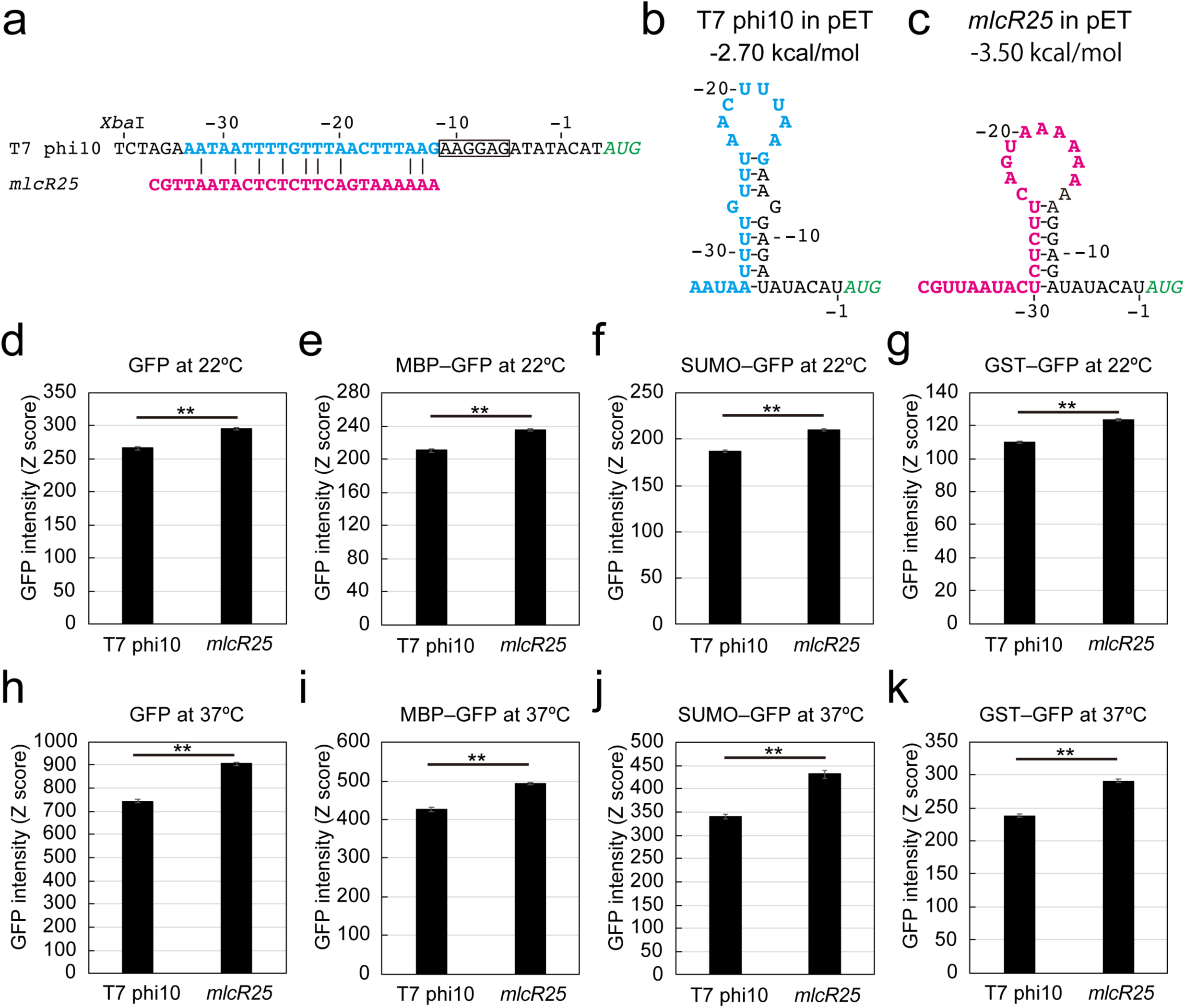
Protein expression in *E. coli* strain BL21(DE3) using T7 expression vectors. (a) Comparison of the T7 phi10 sequence and *mlcR25*. T7 phi10, shown in blue bold, was replaced *mlcR25* shown in magenta bold. The SD sequence is boxed. The initiation codon is shown in green italics. (b, c) Predicted structure of the sequences described in (a). The calculated minimum free energies are indicated. (d-e) Comparison of expression levels of the indicated proteins in cells cultured at 22°C. Fluorescence was measured after IPTG induction for 16 h. (h-k) Comparison of expression levels of the indicated proteins in cells cultured at 37°C. Fluorescence was measured after IPTG induction for 3 h.

## Discussion

This study demonstrated the principle of using TED activity for translation enhancement in *E. coli*. We identified nine genes that have TED activity, among which *mlcR* has the strongest activity. In addition, the activity can be tuned by deletion of part of the sequence. Using TED, we could improve the pET expression system.

We explored sequences with translation enhancement activity by examining various patterns from a preset of *Dictyostelium* gene sequences as a starting material. Alternatively, we may find sequences with a similar effect from randomly designed sequences. The latter approach requires the synthesis of various sequences because there is no template, and thus, is costly and inefficient. From the data currently available in NCBI (https://www.ncbi.nlm.nih.gov/), more than 13,000 coding sequences of *D. discoideum* are annotated. These sequences are promising materials for improving the efficiency of protein production in *E. coli*.

It has been reported that the secondary structure of the sequence upstream of the SD sequence influences the loading and liberation efficiency of the ribosome and subsequent protein expression (de Smit and van Duin 2003; Osterman et al. 2013; Takahashi et al. 2013; Espah Borujeni et al. 2014). By comparing the results of deletion analysis of the *mlcR* sequence with the local secondary RNA structure, we predicted that the sequence that forms a stem loop structure with the SD sequence is effective for TED activity. Based on this criterion, our data empirically indicate that the optimal minimal free energy is calculated to be approximately –3 kcal/mol. This prediction will serve as a basis for finding a better sequence.

We predicted that a sequence forming a moderately stable stem loop upstream of the initiation codon would confer effective TED activity (Fig. 4). This may seem to contradict the view on the relationship between the accessibility of ribosomes to linear mRNA and translation. Interestingly, a recent study reported that the mechanism of internal ribosome-entry site (IRES) works in bacteria as well as in eukaryotes (Colussi et al. 2015). Given that IRES RNAs are known to form intricate structures (Kieft 2008), bacterial ribosomes might recognize and bind to mRNAs having certain structures to initiate translation. Thus, the IRES-like structure may contribute to TED-dependent expression.

An alternative, but not mutually exclusive, mechanism determining the strength of TED activity is mRNA stability (Radhakrishnan and Green 2016). It is known that the features of the 5’-UTR affect the longevity of mRNA (Belasco et al. 1986; Bouvet and Belasco 1992). Furthermore, Komarova et al. (2005) have reported that an A/U-rich 5’-UTR increases mRNA longevity. Unfortunately, due to a technical problem, we could not examine this mechanism in the present study.

Intensive studies have led to a number of improved methods that are associated with potent promoters and optimization of the SD sequence, spacers, and codons (Chen et al. 1994a; Vimberg et al. 2007; Salis et al. 2009; Hanson and Coller 2018). Interestingly, the insertion of a translation-promoting sequence downstream of the initiation codon is also effective (Etchegaray and Inouye 1999; Qing et al. 2003). It is worth emphasizing that TED can be used in combination with all of these methods. One of the advantages of TED is the ease of activity regulation. As shown in Figs. 2A and 3B, even if one promoter is used, various expression levels can be tailored by simple manipulation of the gene sequence, which may be useful for analyzing genes whose phenotype changes depending on the expression level (Ward and Lutkenhaus 1985). Moreover, as the translation mechanism is largely conserved in various bacteria (Laursen et al. 2005), TED is expected to be applicable to expression systems for other bacterial species, such as *Magnetospirillum sp.* (Matsunaga et al. 2007; Uebe and Schüler 2016), *Lactococcus lactis* (Hugenholtz and Smid 2002; Song et al. 2017), and *Bacillus subtilis* (Wong 1995; Song et al. 2015).

In conclusion, tuning of the expression level of a heterologous protein was achieved by using *D. discoideum* gene sequences upstream of the SD sequence. The level of enhancement can be adjusted by the gene used and its length. This method is simple, inexpensive, and easy to perform, and it can be readily adapted for protein expression in both the laboratory and at the industrial scale.

## Acknowledgments

TK was supported by the Japan Society for the Promotion of Science Research Fellowships for Young Scientists.

## Funding

This research was supported by the Japan Society for the Promotion of Science KAKENHI Grant Number 16J08310 to TK.

## Compliance with ethical standards

This article does not contain any studies with human participants or animals performed by any of the authors.

## Conflict of interest

The authors declare that they have no conflict of interest.

