## Supplemental files for "Translation enhancement by a *Dictyostelium* gene sequence in *Escherichia coli*"

**Supplementary Fig. S1 Nucleotide sequence of mlcR–GFP.** (a) Map of *mlcR–gfp*. This fusion was inserted into the pUC19 vector using *Xba*I and *Bam*HI sites. (b) Sequence of *mlcR–gfp*. Colors indicate each fragment shown in (a). The positions of the 5' end of each *mlcR* fragment and the SD sequence-spacer used in Fig. 3 are indicated by white arrows. These fragments were inserted into the vector using the *Xba*I site appended to these ends and the *Bam*HI site appended to the 3' end of *gfp*, as described above.

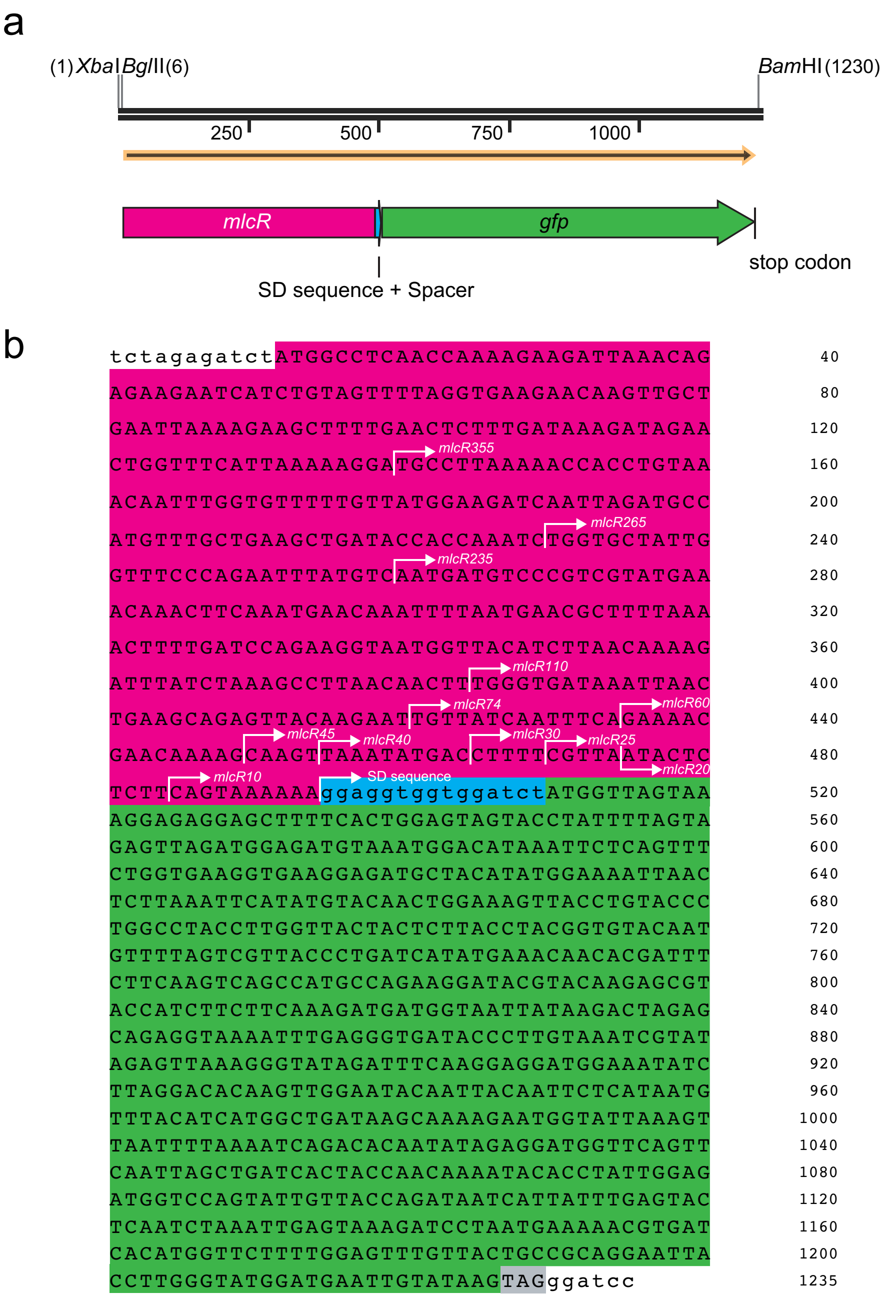

**Supplementary Fig. S2 Growth rate of cells containing mlcR–GFP-pUC19 and the empty vector.** Cells were cultured in LB medium containing 100 µg/ml ampicillin without IPTG at 37°C. The initial cell density was 2.9–3.6 × $10^7$ cells/ml. Error bars indicate the standard deviations (n = 3).

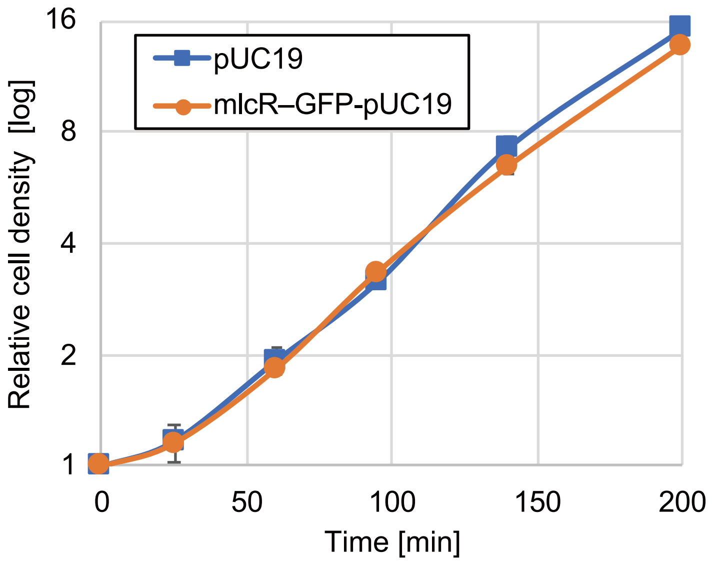

**Supplementary Fig. S3 Comparison of GFP expression levels in strains HST08 and DH5a.** Cells were cultured in LB medium containing 100 μg/ml ampicillin without IPTG at 37°C. Error bars represent the standard errors of the means. n ≥ 170

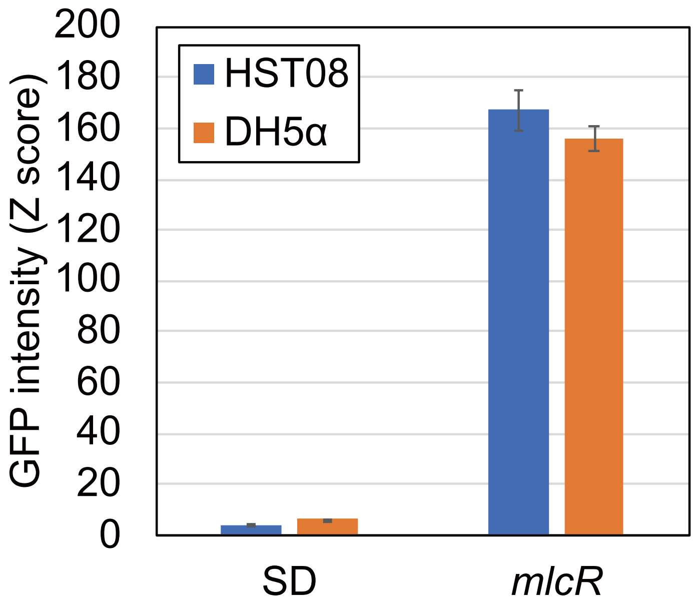

**Table S1. List of genes that used in this study**

| Gene name | Product name | Gene ID in dictyBase[*1] | %AT[*2] | Length [bp][*2] |
|---|---|---|---|---|
| *mlcR* | Myosin regulatory light chain | DDB_G0276077 | 68 | 483 |
| *tubC* | γ-tubulin | DDB_G0271738 | 67 | 1386 |
| *eb1* | EB1 | DDB_G0283607 | 67 | 1518 |
| *tom7* | TOM7 | DDB_G0282897 | 73 | 165 |
| *tom20* | TOM20 | DDB_G0283573 | 63 | 228 |
| DDB_G0288629 | TOM22 | DDB_G0288629 | 71 | 789 |
| *tom40* | TOM40 | DDB_G0277155 | 71 | 942 |
| *tom70* | TOM70 | DDB_G0275389 | 70 | 1605 |
| *H1* | Histone H1 | DDB_G0285319 | 60 | 540 |

*1: http://dictybase.org

*2: not included stop codon
